# Conformational Changes Regulate Metal Coordination in the Catalytic Sites of Cas9

**DOI:** 10.1101/2022.09.10.507422

**Authors:** Anuska Das, Jay Rai, Mitchell O. Roth, Yuerong Shu, Megan L. Medina, MacKenzie R. Barakat, Hong Li

## Abstract

The control of CRISPR-Cas9 activities plays an essential role in its safe adoption for clinical and research applications. Previous studies have identified conformational dynamics of Cas9 as a key regulatory element for its enzymatic activity. The molecular basis for how conformations influence the catalytic processes at both nuclease domains, however, remains elusive. Here we present well-resolved cryo-EM structures of the active *Acidothermus cellulolyticus* Cas9 (AceCas9) complexes representing pre-cleavage (2.9 Å), two reaction intermediate (2.2 Å and 2.4 Å) and two post-cleavage (2.6 Å and 2.7 Å) states that reveal active site rearrangement during DNA binding and phosphodiester bond breakage by Cas9. Strikingly, large domain rearrangements in Cas9 triggered by the cognate DNA result in subtle changes in active sites that facilitate coordination of metal ions required for catalysis. The two reaction intermediate structures further reveal small oscillations in domain conformations, which alternates the reactive states of the two catalytic centers, thereby coupling cleavage of the two DNA strands. Consistent with the roles of conformations in organizing the active sites, adjustments of metal coordination ligands lead to altered metal specificity.

## Introduction

The use of the Clustered Regularly Interspaced Short Palindromic Repeat (CRISPR) and associated protein 9 (Cas9) has revolutionized genome engineering technology^1,2^. Efforts in understanding the underlying molecular basis for its catalytic efficiency and specificity can greatly improve the utility of Cas9. The Cas9 endonuclease targets DNA that contains a 20-24 nucleotide (nt) complementary region (protospacer) to the associated CRISPR RNA (crRNA) and a unique 3-8 nt segment downstream of the protospacer (PAM, Protospacer Adjacent Motif). A hallmark of Cas9 function is the DNA-triggered conformational changes that control the catalytic efficiency and substrate specificity^3^. Extensive structural and biophysical characterizations have revealed that two conserved lobes, the nucleic acid binding lobe (REC) and the nuclease lobe (NUC), each comprising half of the protein, undergo concerted motions^3,4^ The REC lobe is primarily responsible for anchoring the conserved repeat and anti-repeat helix between the crRNA and the tracrRNA (or the tetraloop-linked helix in the single-guide RNA, sgRNA) and the guide-DNA heteroduplex. Part of the REC lobe buttresses the guide-DNA heteroduplex as it enters the cleavage position and disengages after the cleavage. The NUC lobe includes two catalytic domains: RuvC and HNH, for cleaving the nontarget (NTS) and the target DNA (TS) strand, respectively. Concomitant with the engagement of the REC lobe with the guide-DNA heteroduplex, the HNH domain undergoes a large conformational change to be docked onto the target strand^5,6^. Mutations in REC, the RuvC-HNH linker or the guide-DNA heteroduplex that are known to change catalytic efficiencies also impact the rate of HNH conformational placement^5,7–9^. Correctly placed HNH, but not necessarily its catalytic activity, activates the RuvC domain for the cleavage of the nontarget DNA strand^10^. Similar mutations impact the cleavage efficiency of both the target and nontarget strand to the same degree^5^, suggesting a coordination between the HNH and RuvC domains.

Divalent metal ions are required for catalysis mediated by both the HNH and the RuvC domain^11,12^, and interestingly, for the conformational change of the HNH domain^8^. The metal coordination environment in Cas9 has rarely been observed often due to the use of non-reactive conditions or to low structure resolution. Though metal coordination at the catalytic centers is expected based on structural studies of homologous domains in other DNA cleaving enzymes, how it is linked to Cas9 dynamics, however, is completely unknown. The HNH domain has the typical ßßα-metal fold and contains the key Asn (on a), as well as Asp and His (on ß1) residues^13^. The RuvC domain of Cas9 contains a six-stranded mixed ß-sheet (541236, ↑↑↑↓↓↓) sandwiched by several helices and the key Asp (on ß1), Glu (on ß4), and His (on aA) residues. While a single divalent ion is required for the HNH-mediated target strand cleavage, two are required for the RuvC-mediated nontarget strand cleavage^10,13^. Metal ions, especially Mg^2+^, are extremely sensitive to their coordination ligands, which form the chemical basis for nucleic acid enzymes to confer substrate specificity^14^ Direct observation of the intact active sites of Cas9 including metal ion holds the key to understanding and improving Cas9 cleavage chemistry.

### Acidothermus cellulolyticus

Cas9 (AceCas9) belongs to the Type II-C subtype that are, generally, smaller in size but contain the identical catalytic residues to those in the commonly used Type II-A Cas9 such as SpyCas9. In addition, the RuvC domain used by AceCas9 is also the sole catalytic domain of another commonly used CRISPR endonuclease, Cas12, such as Cpf1^15^. AceCas9 is moderately thermophilic and specific to DNA protospacers associated with a 5’-NNNCC-3’ PAM. Though the wild-type AceCas9 has slower rate of cleavage than that of SpyCas9, protein directed engineering has resulted in a catalytic enhanced AceCas9 that has a comparable cleavage rate to that of SpyCas9^9^. Importantly, AceCas9 is uniquely sensitive to DNA methylation and does not cleave DNA with a 5’-NNN^m^CC-3’ PAM^16^. Both the unique and the conserved biochemical properties of AceCas9 make it an excellent candidate as a biotechnology tool and a model system for studying genome editing enzymes. Here, we assembled the intact AceCas9 with a cognate DNA substrate under a reactive condition and studied the structures of AceCas9 complexes populated in the reaction mixture by electron cryomicroscopy (cryo-EM). We captured five distinct states of AceCas9 at 2.2 – 2.9 Å resolutions including two catalytic intermediates (Supplementary Figure 1 & Supplementary Table 1). The well-resolved density reveals detailed coordination chemistry of the metal ions in the HNH and RuvC catalytic sites and how they are regulated by conformational changes.

## Results

### Domain Movements during Catalysis

The recombinantly expressed full-length AceCas9, assembled with an *in vitro* transcribed sgRNA, was first tested for its dependence on metal ions using individually labeled DNA substrates (Figure 1). While the target strand cleavage by the HNH domain is supported by Mg^2+^, Mn^2+^, and Co^2+^, the nontarget strand cleavage by the RuvC domain is supported by Mg^2+^ and Mn^2+^, suggesting that RuvC has more stringent requirement for metal coordination properties than HNH.

**Figure 1.**
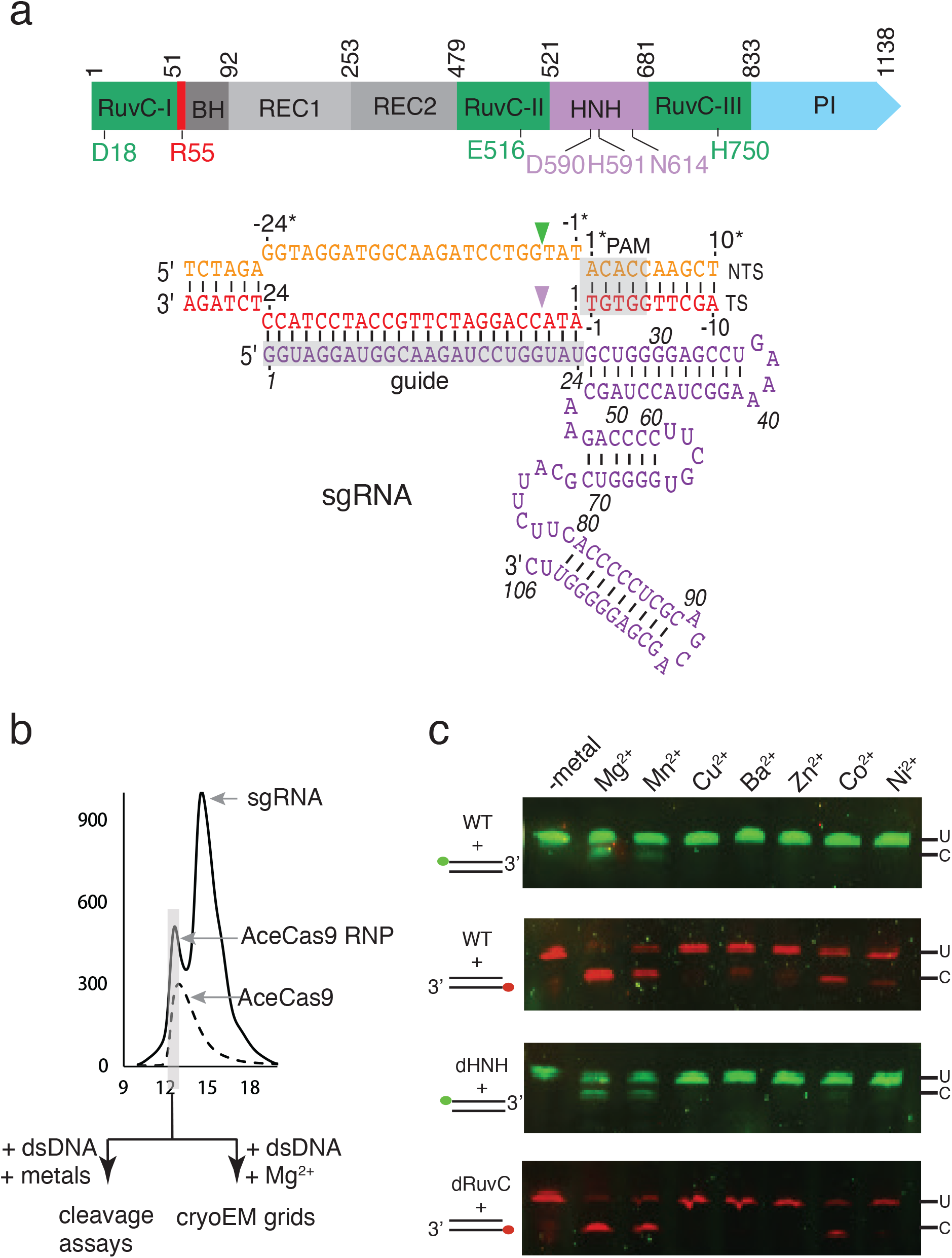
Features of the *Acidothermus cellulolyticus* Cas9 (AceCas9) system and dependency on metal ions. (a) Top: Domain organization of AceCas9. Regions in AceCas9 corresponding to structural domains are colored and labeled. Relevant residues are labeled. Bottom: Schematic diagram of nucleic acids used in this study with nucleotide numbers labeled. Cleavage sites for the nontarget (NTS) by the RuvC and the target strand (TS) by the HNH domain are indicated by a green and a pink triangle, respectively. The Protospacer Adjacent Motif (PAM) is highlighted in gray. (b) Overlay of the gel filtration profiles of AceCas9 protein and its ribonucleoprotein (RNP) complex assembled with the single-guide RNA (sgRNA) shown in (a). Samples collected for biochemistry and cryoEM analysis are highlighted by a gray box. (c) Cleavage results of dsDNA assembled with either HEX labeled TS (red) or 6-FAM labeled NTS oligo (green) at 10 nM by AceCas9 or its catalytic mutants at 1 μM in presence of various divalent ions at 10 mM.

To capture the structures of AceCas9 in presence of catalytic metal ions, we assembled an active AceCas9-sgRNA-DNA complex in a buffer containing Mg^2+^ and allowed the reaction to proceed for 15-30 minutes before preparation of cryo-EM samples (Figure 1). Particles from two independent data sets were sorted and classified extensively (Supplementary Figures 1 & 2, Supplementary Table 1), which resulted in six different classes based on the location (or absence) of the HNH and the REC2 domains. The excellent quality of density around the DNA cleavage sites in all structures, especially that of the target strand, allowed the classes to be sorted into the structures of pre-cleavage (state A), cleavage intermediate (states B1 and B2), post-cleavage (states C1 and C2), and target strand bound state (state D) (Figure 2).

**Figure 2.**
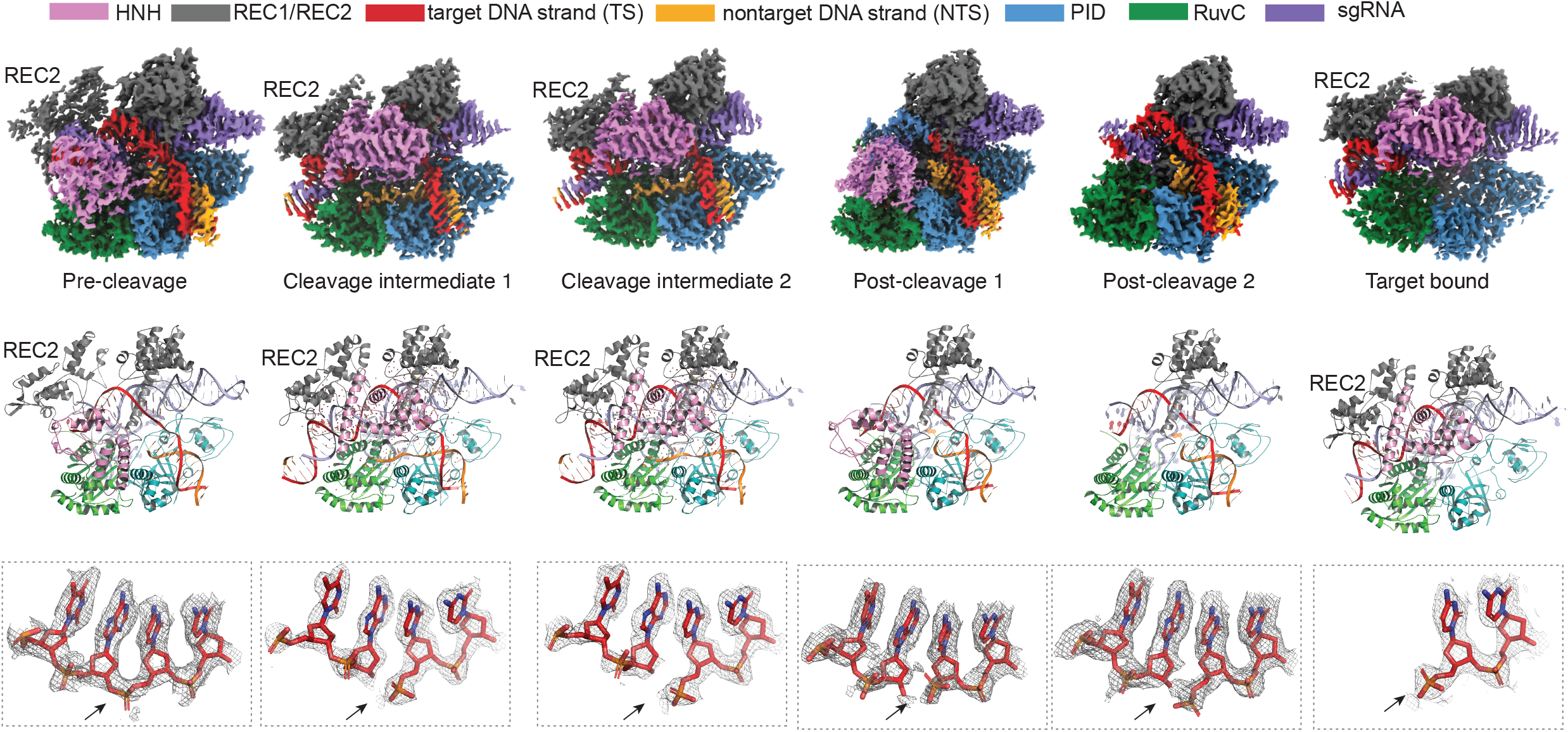
The observed Cryo-EM structures at six cleavage states. Top, electron potential density maps corresponding to the pre-cleavage (A), the cleavage intermediates (B1 & B2), the post-cleavage (C1 & C2), and the target (only) bound state. Middle, cartoon representations of the structural models corresponding to the maps at top. Bottom, closeup views of the cleavage site of the TS overlaid with the density for each of the corresponding state on top and middle. The arrow indicates the position of the scissile phosphate bond.

The most significant structural difference among the various states is the location of the HNH and the REC2 domain. Both the pre- and post-cleavage states (A, C1 and C2) (OPEN) are characterized by HNH either being placed away from the target cleavage site or disordered. In these states, the REC2 domain is also loosely associated with the guide-target heteroduplex or disordered. By contrast, the other three states (B1, B2 and D) (CLOSED) form a compact structure with both HNH and REC2 domains closely engaging the DNA substrate (Figure 2). To reach the CLOSED from the OPEN conformation, the HNH domain rotates 180 degrees and the REC2 domain swings towards the RuvC domain (Supplementary Figure 3). Surprisingly, the two reaction intermediate states, though both are CLOSED and well ordered, exhibit subtle but clear domain differences (Supplement Figure 3). We mutated the residues lining the interface between the guide-target heteroduplex and the RuvC domain and observed severe impacts on *in vivo* DNA cleavage activity (Supplementary Figure 4). The results support the importance of the HNH/REC1-mediated dynamic interactions in reaching the CLOSED conformation.

### A Type II-C Cas9-specific R-loop Lock

Regardless of the state, the R-loop structure is correctly formed with the target strand base paired with the spacer region of the sgRNA (Figure 2). However, an important difference in R-loop structure is observed between the CLOSED and the OPEN conformation. In the CLOSED conformation, a Type II-C Cas9-specific loop of ~12 amino acids immediately before the Bridge Helix (BH) advances towards the forked DNA to lock the R-loop for DNA cleavage (R-loop lock) (Figure 3). Residue Arg55 engages the last TS-NTS base pair, T(-1)-A(1*), through a cation-π interaction, resembling a “tongue depressor”, allowing the upstream NTS nucleotides to feed into the RuvC active site. In the OPEN conformation, however, T(-1*) of the NTS returns to be stacked on A(1*) with Arg55 stacks on its base, also through the cation-π interaction (Figure 3), thereby directing NTS away from the RuvC center. Mutation of Arg55 to Trp or Tyr largely retained while that of Arg55 to Ala abolished the activity (Figure 3). The conserved presence of the R-loop lock, especially Arg55, in Type II-C Cas9 may be a molecular strategy to compensate for their relatively weak DNA unwinding activity^17^.

**Figure 3.**
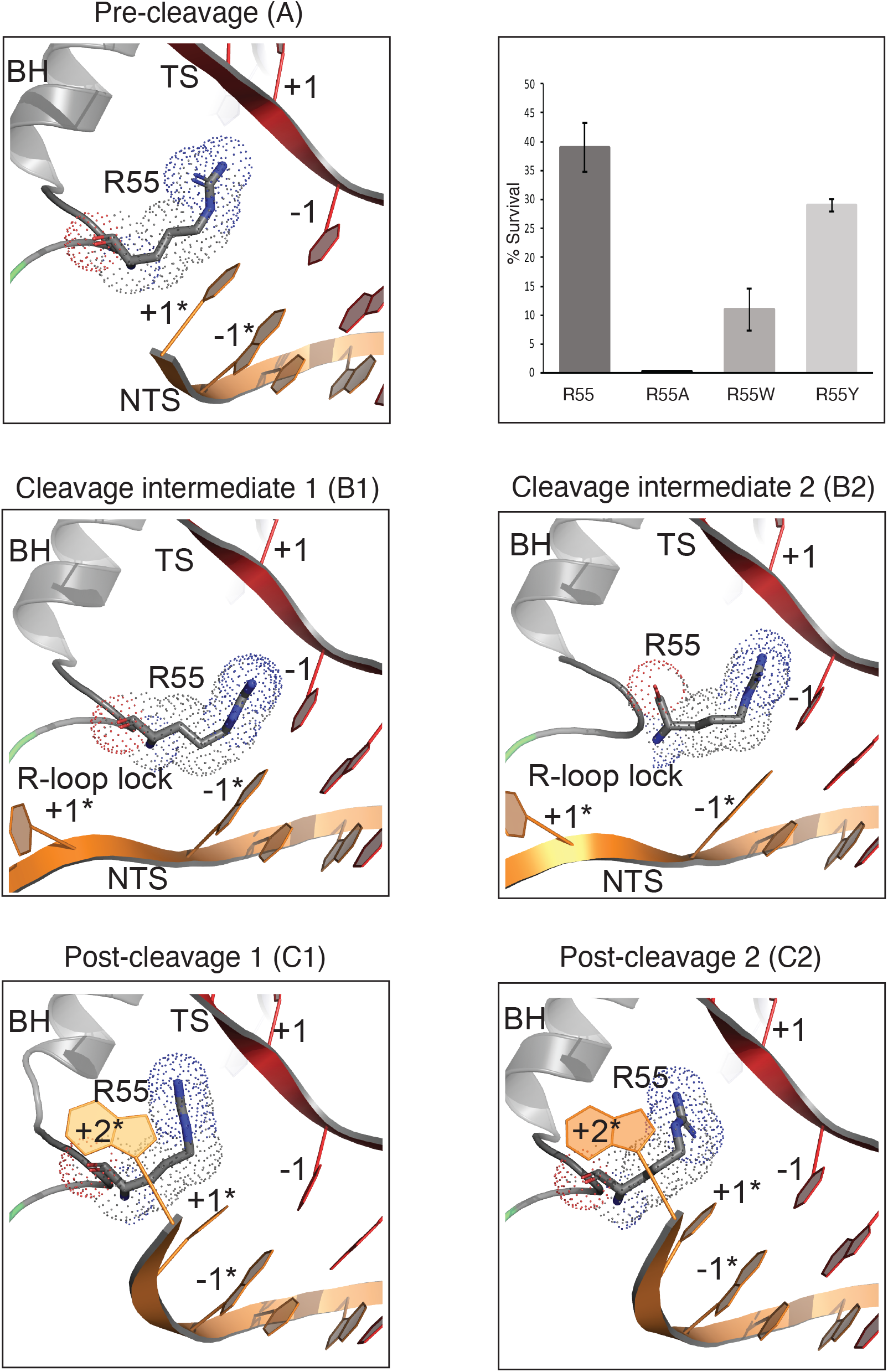
R-loop lock structure in different functional states and functional assay results. The R-loop lock residue Arg55 is shown in stick model with dotted van der Waals surface at the fork formed between the NTS and the TS DNA in R-loop. Each functional state is labeled as in Figure 2. Top right, calculated survival rates of the wild-type (R55) and Arg55 mutations (R55A, R55W, R55Y) shown in bar diagram. The survival rate is calculated as the ratio of the colonies on the arabinose-chloramphenicol plates to that of chloramphenicol only plates. The experiment was performed in triplicate and plotted with error bars showing the standard deviations.

### Dependence of Metal Coordination on Domain Movements

In the three OPEN state (A, C1 and C2), we did not observe convincing density that can account for the coordinated metal ions in either active site. This is in part due to the absence of DNA substrate in the enzyme active sites that is known to play a critical role in coordinating metals^14^. In the two CLOSED states (B1 and B2), however, both active sites closely engage the two respective scissile phosphate groups and contain well-coordinated metal ions. Unexpectedly, the two states reveal subtle but clear differences in active site geometries linked to their different domain conformations.

The HNH catalytic center contains a single metal as expected that has a near perfect octahedral coordination with two water molecules, the carboxylate of Asp590, the carboxyamide of Asn614, and a non-bridging oxygen of the scissile phosphate (Figure 4). This configuration is consistent with a mechanism where the metal ion both prepares the scissile bond for the nucleophilic attack and stabilizes the negatively charged pentavalent intermediate. The nearby His591 does not coordinate with the metal and acts as the general base to deprotonate the nucleophile water. It also is consistent with the fact that other octahedron-coordination metals can support the cleavage reaction (Figure 1). Strikingly, the HNH active site configuration differs in the position of the leaving 3’ OH group between the two reaction intermediates. In B1 where the HNH domain forms a tighter grip on the target strand, the leaving 3’ OH remains tethered to the metal at 2.7 Å while in B2 where the HNH undocks slightly (Supplementary Figure 3), it pulls away from the metal by additional 0.9 Å with the loss of connecting density (Figure 4a), indicating that B2 follows B1 on the HNH reaction path.

**Figure 4.**
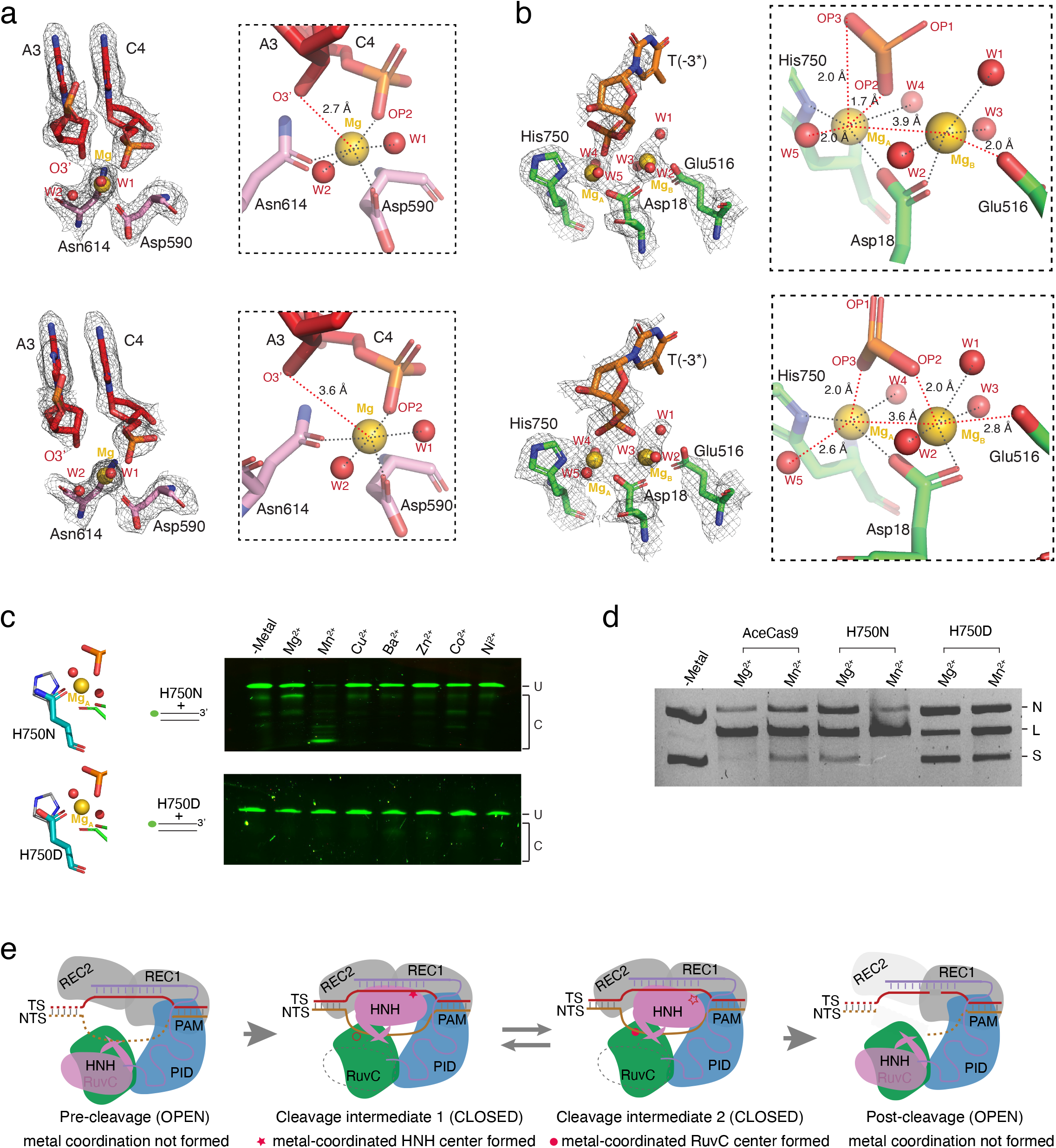
The observed density and geometries at the HNH and the RuvC catalytic centers in the reaction intermediates. Density is shown in gray mesh. Protein residues and DNA nucleotides are shown as colored stick models. Metal ions are shown as gold spheres. Water molecules are shown as red spheres. (a) Density and structures of the HNH catalytic center of the intermediate B1 (top) and B2 (bottom). Insets shows close-up views of the metal coordination environment. Black dash lines indicate coordination distances between 1.7 Å – 2.1 Å. Red dash lines indicate the distances varied between the B1 and B2 intermediates. (b) Density and structures of the RuvC catalytic center of the intermediate B1 (top) and B2 (bottom). Insets shows close-up views of the metal coordination environment. Black dash lines indicate coordination distances between 1.7 Å – 2.1 Å. Red dash lines indicate the distances varied between the B1 and B2 intermediates. (c) In vitro DNA cleavage activities on double stranded DNA oligo substrate by the RuvC domain of AceCas9 mutants His750Asn (H750N) and His750Asp (H750D) in presence of various of divalent ions. The nontarget strand is labeled with 6-FAM (Fluorescein) fluorescence probe (green dot) at its 5’ end. The stick models represent putative structures of the mutated residues (teal) in comparison with the wild-type His750 (gray). “U” denotes uncleaved and “C” denotes cleaved DNA substrate. (d) Comparison of DNA plasmid cleavage activities of AceCas9, H750N and H750D in presence of Mg^2+^ or Mn^2+^ ions. “N” denotes nicked, “L” denotes linearized, and “S” denotes supercoiled plasmid DNA. 500 nM of RNP was used and the reactions were incubated for 15 minutes at 50°C.(e) Proposed active site formation in AceCas9 through domain movements and the coupling between two active sites. The RuvC and HNH active sites are not formed with coordinated metals in the pre- or post-cleavage state but are formed in the cleavage intermediates. However, the two active sites alternate between reactive (solid red) and non-reactive (open red) in the two cleavage intermediates. Domains are colored consistently as those in Figure 2.

Interestingly, the RuvC center shows an opposite reaction path than that of the HNH center with respect to two intermediate conformations. The RuvC center contains two well-coordinated Mg^2+^ that also show a different arrangement between the two reaction intermediates (Figure 4b). In B1, metal A is coordinated with two water molecules, the carboxylate of Asp18, Nδ1 of His750, and a non-bridging oxygen of the scissile phosphate in a tetrahedral geometry. While in B2 where the HNH domain undocks slightly, the nonbridging oxygen of the scissile phosphate of the nontarget strand repositions and coordinates with metal B instead of A (Figure 4b). Concomitantly, metal B that is coordinated in a perfect octahedral geometry with three water molecules, the carboxylates of both Asp18 and Glu516 in B1, loses its coordination with Glu516 in B2 while gaining that with a non-bridging oxygen of the scissile phosphate (Figure 4b). As a result, the two metal ions shift from being separated by 3.9 Å in B1 to 3.6 Å in B2, a distance believed to be important for initiation of catalysis^14^. The observed geometry and the density connectivity indicate that, unlike the HNH center, the intermediate B2 is the state immediately after the nucleophilic attack and precedes B1 on the RuvC reaction pathway.

The metal coordinating Glu516 is on the well-conserved ß4 of the RuvC domain that is linked directly to the hinge motion of the HNH domain (Supplementary Figure 3). In B2 where HNH is relaxed after cleavage, Glu516 retracts slightly to allow metal B to move closer to metal A and to coordinate with the non-bridging oxygen, which helps to bring the nucleophile in place to activate RuvC cleavage. When HNH returns to B1, Glu516 is now poised to coordinate with metal B, which separates the two metals 3.9 Å apart and returns RuvC to a pre-cleavage state. The change in metal coordination in the RuvC catalytic center resulting from the subtle rotation of HNH is consistent with the observation that HNH leads RuvC in cleavage^10,18^ and suggests a linked catalytic processes between the two active sites.

The conformation-mediated Glu516 repositioning is significantly more pronounced when AceCas9 transitions from the CLOSED to the OPEN conformation. In the OPEN conformation and regardless of if HNH is ordered, the ß4 strand of RuvC stays tightly parallel with ß1, which makes Glu516 sidechain point away from metal B (Supplementary Figure 3). In this conformation, RuvC remains inactive even if DNA is placed in its active site. The geometric property and strategic location of Glu516 mediate the control of RuvC activity through HNH conformations.

### Tuning Metal Specificity of AceCas9

Given the tight control of the active centers through AceCas9 conformations, we wonder whether its metal specificity may be altered. We mutated His750 (His750 to Asn, H750N and Hs750 to Asp, H750D) that coordinates with metal A and tested the metal dependency of the mutants. Unlike the wild-type AceCas9 that prefers Mg^2+^, The H750N variant prefers Mn^2+^ (Figure 4c, Figure 4d & Supplementary Figure 6). In addition, H750N showed enhanced 3’-exonuclease activity based on the shortened products from the Mn^2+^-supported cleavage (Figure 4c, Figure 4d & Supplementary Figure 6). Interestingly, although H750D mimics the active center of Cas12 RuvC domain, it showed no preference for and significantly reduced DNA cleavage activity with either Mg^2+^ or Mn^2+^ (Figure 4c, Figure 4d & Supplementary Figure 6). No collateral DNase activity is observed in H750D either (Supplementary Figure 6), unlike Cas12, supporting a tighter control of the RuvC domain in Cas9 than in Cas12. H750N may be utilized in applications where Mn^2+^ concentration is elevated while that of Mg^2+^ is reduced.

## Discussion

The series of functional complex structures of the methylation-sensitive Cas9, AceCas9 at near atomic resolutions reveal both large and subtle conformation changes necessary for catalysis (Figure 4e). We show that the catalytically relevant binding of metal ions to AceCas9 only occurs in presence of the substrate as well as a correct the enzyme conformation. Two on-path reactive states were observed for the RuvC and the HNH catalytic center, respectively, in the two cleavage intermediate states that differ slightly in conformation. Most remarkably, the reaction paths of the two catalytic centers are opposite to each other, suggesting a coordination between them linked by conformations. In sequence-specific DNase such as restriction endonucleases, metal coordination is often sensitive to the cognate base pairs^14^. In Cas9 where sequence specificity is not required, however, metal coordination is exploited by enzyme conformational changes in control of its activity. As catalytic efficiency seems to be correlated with the canonical configuration of the metal ions, slight changes in coordination environment such as those between the two cleavage intermediates of AceCas9, or those due to guide-target mismatches, protein mutations, or other compositional changes, can thus tune activities through metal coordination.

In two-metal catalyzed phosphoryl transfer reactions, metal B is the perfect target for regulation, as it is known to be more critical to catalysis than metal A^14^. The universally conserved glutamate (Glu516 in AceCas9) offers a long sidechain-tethered carboxylate and is exploited by Cas9 to regulate RuvC activity. Subtle changes in HNH domain are observed to influence Glu516-mediated coordination to metal B. Metal A often has imperfect coordination geometry and can be replaced by other catalytic groups such as by an amino acid general base. Consistently, when the HNH site is superimposed with the RuvC site, the HNH-associated Mg^2+^ more closely mimics metal B (Supplementary Figure 5). Significantly, we are able to alter the coordination of metal B by mutating His750 to asparagine, which results in an AceCas9 variant that favors Mn^2+^.

The catalytic residues in both HNH and RuvC are universally conserved in all Cas9 known so far and they also bear a reasonable overall conservation (Supplementary Figure 7). The observed coordination between the two active sites in Acecas9 likely apply to all Cas9 in which the catalytic events of the two active sites are synced through the oscillating reaction intermediate states. Indeed, careful enzyme kinetic studies of SpyCas9 showed comparable rates of cleavage by HNH and RuvC^10^, and interestingly, that HNH conformation regulates RuvC cleavage activity^18^. The linked catalysis provides a means to ensure coordinated cleavage of both DNA strands by two separate active sites.

Unlike Cas9, the RuvC domain in Cas12 is required to cleave both the target and the nontarget strands. In addition, several characterized Cas12 enzymes show strong single-stranded DNA cutting following activation by the cognate dsDNA, an activity apparently lacking in Cas9^19^. Though the AceCas9 His750Asp variant mimics the active center and the metal dependency of Cas12, it has significantly reduced dsDNA activity and remains free of ssDNA cleavage activity. The mechanism for the constitutively activated RuvC in Cas12 remains to be fully elucidated. However, the lack of an analogous HNH domain-mediated control may explain the different RuvC activity between Cas9 and Cas12.

## Materials and Methods

### Protein and nucleic acid sample preparation

AceCas9 was cloned in pET28b vector and expressed in *Escherichia coli* BL21 strain, and was purified as previously described^16,20^. Briefly, the protein was purified by Nickel affinity chromatography followed by heparin ion exchange. The heparin purified protein was stored at −80°C for further use. The wild type and mutant proteins used for *in vitro* assays were further purified with size exclusion chromatography and stored at a 10μM concentration. All of the mutations were introduced using the Q5 site directed mutagenesis kit (New England Biolabs, Ipswich, MA) with primers listed (Supplementary Table 2).

The 94 nt sgRNA was in vitro transcribed with T7 RNA polymerase (Supplementary Table 2) and purified by phenol-chloroform extraction followed by gel filtration. The DNA used in cryoEM and biochemical assays were purchased from Eurofins Genomics (Louisville, KY). The heparin purified full-length protein was mixed with sgRNA at a 1:2 molar ratio and the RNP was purified via size exclusion chromatography using Superdex 200 increase column (Figure 1). The AceCas9-RNA-DNA ternary complex was reconstituted by adding preannealed dsDNA substrate DNA to the RNP at a 1:1 molar ratio. Metal ions were made to a final concentration of 40mM to the complex to initiate the nuclease activity. Magnesium chloride was used for the complex used in cryoEM sample preparation.

### Cryo-EM sample preparation, data collection and 3D reconstruction

The Cas9-RNA-DNA ternary complex at approximately 2.0 absorption units at 260 nm (A_260_) and in 4μl volume was applied to glow-discharged UltrAuFoil 300 mesh R1.2/1.3 grids (Quantifoil). The sample was let to absorb for 10 seconds on girds followed by 2 seconds blotting at 19°C and 100% humidity. The grids were plunged freeze in liquid ethane cooled in liquid nitrogen using FEI Vitrobot Mark IV.

Two data sets were collected at the Laboratory for BioMolecular Structure (LBMS) of the Brookhaven National Laboratory on a Krios G3i cryo transmission electron microscope equipped with Gatan K3 direct electron detector (ThermoFisher Scientific). All the images were collected at 105000x magnification with a corrected pixel size of 0.412 Å/pixel and −1 to −2.2 μm defocus range in a counted super-resolution mode with 20 eV energy filter. Motion correction was carried out in bin 2 using MotionCorr2^21^ and contrast transfer function (CTF) estimation was performed with Gctf^22^. LoG-based auto-picking algorithm in RELION-4.0 was used for particle picking. cryoSPARC^23^ was used for template-based particle picking and initial 2D classification to eliminate bad particles. The gold-standard Fourier Shell Correlation (FSC) at a value of 0.143 was used for resolution estimation. Local resolution was estimated using RELION-4.0.

A total of 13,653 micrographs were collected from two independent data sets and 12,757 were selected for further processing. Several rounds of 2D classification were executed to result in 5,817,494 good particles for 3D classification in RELION-4.0. The classes with similar features, especially with respect to the HNH domain and REC2 domain, were combined and refined using customized 3D masks. Particles contributed to the class with HNH and REC2 domains closed was further classified with 3D custom masks leading to two distinct subclasses with improved resolution. Multiple rounds of CTF refinement were carried out to reach the final 3D reconstructions (Supplementary Figures 1 & 2). Structural models were built in COOT^24^ and refined in PHENIX^25^ to satisfactory stereochemistry and real space map correlation parameters (Supplementary Table 1).

### In vitro oligo cleavage assay

The wild-type or mutant AceCas9 was pre-incubated at 37°C for 30 minutes with sgRNA at 1:1 molar ratio to form the RNP. The fluorescein amidites (FAM) labeled NTS with molar excess of non-labeled TS or the hexachlorofluorescein (HEX) labeled TS with molar excess of non-labeled NTS was annealed by heating at 95°C for 5 minutes and cooled slowly to room temperature. The pre-annealed dsDNA substrate at 10 nM of the labeled strand was added to the RNP at a final concentration of 500 nM – 1μM along with divalent metal ions at 10 mM. The reactions were incubated at 50°C for a predetermined time before being quenched with the addition of an EDTA (Ethylenediamine tetraacetic acid)-containing stop buffer. The oligo products were resolved on an 8M urea 15% polyacrylamide denaturing gel and visualized on Bio-Rad ChemiDoc gel imaging system with 488 nm and 580 nm excitation wavelength for the FAM and the HEX labeled DNA, respectively.

### In-vitro plasmid cleavage assay

The wild type or mutant AceCas9 was mixed with sgRNA at 1:1 ratio and incubated at 37°C for 30 minutes to form RNP. For metal dependent cleavage assay, different divalent metals at 5-10 mM concentration and target plasmid at 6 nM were added to RNP at 500 nM – 1μM concentration and incubated for different periods of time. The reactions were quenched by adding 5x stop buffer (25 mM Tris pH 7.5, 250 mM EDTA pH 8.0, 1% SDS, 0.05% w/v bromophenol blue, 30% glycerol). The products were resolved on 1% agarose gel and stained with ethidium bromide. For M13 ssDNA collateral cleavage assay, M13 ssDNA plasmid at 2 μM concentration was added to the plasmid cleavage reaction and incubated for 50 minutes at 37°C. The reactions were stopped and resolved on a 0.7% agarose gel and stained with ethidium bromide. The gels were visualized by Bio-Rad Chemidoc gel imaging system.

### Cellular survival assay

Cell survival assay was performed as previously described^26^ with two modifications. The DNA sequence targeted in the p11-LacY-wtx1 encoding ccdB toxin is flanked by the minor PAM sequence 5’-NNNAC-3’ (p11-LacY-wtx1-AC) for AceCas9. The AceCas9-encoding plasmid contains the previously characterized Glu839Arg and Glu840Tyr mutation (RY)^27^ (AceCas9-RY) that enables survival in the cell containing the minor 5’-NNNAC-3’ PAM and a graded survival activity. Additional AceCas9 mutations were constructed on the RY background. To perform survival assay, *E.coli* BW25141 (gift from D.Edgell) containing the p11-LacY-wtx1-AC were transformed with 100-200ng of plasmids encoding the AceCas9-RY variants. The cells were recovered in super-optimal broth (SOC) at 37°C for 30 min, followed by induction with 0.05 mM Isopropyl *β*-D-1-thiogalactopyranoside (IPTG) and recovered for additional 1 hour. The cells were then plated in chloramphenicol (52 μg/ml) or chloramphenicol-arabinose (10mM) containing plates. The plates were incubated at 37°C for 16-20 hours then colony forming units (CFU) were counted on each plate. Survival percentages were calculated as the ratio of CFU on the chloramphenicol-arabinose plates to that of the chloramphenicol only plates. The percentages with errors from three technical replicates were plotted in Microsoft excel.

## Data Availability

The atomic coordinates and associated density maps have deposited at Protein Data Bank with accession codes 8D2N (Pre-cleavage), 8D2L (Cleavage intermediate 1), 8D2K (cleavage intermediate 2), 8D2Q (Post-cleavage 1), 8D2O (Post-cleavage 2) and 8D2P (Target bound) and EMBD 27143 (Pre-cleavage), 27142 (Cleavage intermediate 1), 27141 (Cleavage intermediate 2), 27146 (Post-cleavage 1), 27144 (Post-cleavage 2) and 27145 (Target bound).

## Acknowledgments

This work was supported by NIH grant R01 GM101343 to H.L. The authors also acknowledge the use of instruments at the Biological Science Imaging Resource supported by Florida State University. The Titan was funded from NIH grant S10 RR025080. The BioQuantum/K3 was funded from NIH grant U24 GM116788. The Vitrobot Mk IV was funded from NIH grant S10 RR024564. The Solaris Plasma Cleaner was funded from NIH grant S10 RR024564. The DE-64 was funded from NIH grant U24 GM116788. The Laboratory for BioMolecular Structure (LBMS) is supported by the DOE Office of Biological and Environmental Research (KP160711).

## Author Contributions

A.D. and H.L. designed the experiments. A.D. purified the samples. J.R. and A.D. prepared cryoEM grids and collected data. J.R. performed the cryoEM analysis. M.R., M.M. and M.B. made the mutants and performed the cell survival assays. A.D. performed the *in-vitro* assays with the assistance of Y.S.. A.D. J. R. and H.L analyzed data, made the figures, and wrote the manuscript. All edited manuscript.

## References

1. Hsu, P.D., Lander, E.S. & Zhang, F. Development and Applications of CRISPR-Cas9 for Genome Engineering. Cell 157, 1262–1278 (2014).

2. Sternberg, S.H. & Doudna, J.A. Expanding the Biologist’s Toolkit with CRISPR-Cas9. Mol Cell 58, 568–574 (2015).

3. Jiang, F. & Doudna, J.A. CRISPR-Cas9 Structures and Mechanisms. Annu Rev Biophys 46, 505–529 (2017).

4. Jiang, F. et al. Structures of a CRISPR-Cas9 R-loop complex primed for DNA cleavage. Science 351, 867–71 (2016).

5. Sun, W. et al. Structures of Neisseria meningitidis Cas9 Complexes in Catalytically Poised and Anti-CRISPR-Inhibited States. Mol Cell 76, 938–952 e5 (2019).

6. Zhu, X. et al. Cryo-EM structures reveal coordinated domain motions that govern DNA cleavage by Cas9. Nat Struct Mol Biol 26, 679–685 (2019).

7. Chen, J.S. et al. Enhanced proofreading governs CRISPR-Cas9 targeting accuracy. Nature 550, 407–410 (2017).

8. Dagdas, Y.S., Chen, J.S., Sternberg, S.H., Doudna, J.A. & Yildiz, A. Conformational Dynamics of Cas9 during DNA Binding. Biophysical Journal 112, 71a–71a (2017).

9. Hand, T.H. et al. Catalytically Enhanced Cas9 through Directed Protein Evolution. CRISPR J 4, 223–232 (2021).

10. Gong, S.Z., Yu, H.H., Johnson, K.A. & Taylor, D.W. DNA Unwinding is the Primary Determinant of CRISPR-Cas9 Specificity. Biophysical Journal 114, 251a–251a (2018).

11. Jinek, M. et al. A programmable dual-RNA-guided DNA endonuclease in adaptive bacterial immunity. Science 337, 816–21 (2012).

12. Gasiunas, G., Barrangou, R., Horvath, P. & Siksnys, V. Cas9-crRNA ribonucleoprotein complex mediates specific DNA cleavage for adaptive immunity in bacteria. Proc Natl Acad Sci U S A 109, E2579–86 (2012).

13. Yang, W. Nucleases: diversity of structure, function and mechanism. Q Rev Biophys 44, 1–93 (2011).

14. Yang, W., Lee, J.Y. & Nowotny, M. Making and breaking nucleic acids: two-Mg2+-ion catalysis and substrate specificity. Mol Cell 22, 5–13 (2006).

15. Swarts, D.C. & Jinek, M. Cas9 versus Cas12a/Cpf1: Structure-function comparisons and implications for genome editing. Wiley Interdiscip Rev RNA 9, e1481 (2018).

16. Das, A. et al. The molecular basis for recognition of 5’-NNNCC-3’ PAM and its methylation state by Acidothermus cellulolyticus Cas9. Nat Commun 11, 6346 (2020).

17. Ma, E., Harrington, L.B., O’Connell, M.R., Zhou, K. & Doudna, J.A. SingleStranded DNA Cleavage by Divergent CRISPR-Cas9 Enzymes. Mol Cell 60, 398–407 (2015).

18. Raper, A.T., Stephenson, A.A. & Suo, Z. Functional Insights Revealed by the Kinetic Mechanism of CRISPR/Cas9. J Am Chem Soc 140, 2971–2984 (2018).

19. Chen, J.S. et al. CRISPR-Cas12a target binding unleashes indiscriminate single-stranded DNase activity. Science 360, 436–439 (2018).

20. Tsui, T.K.M., Hand, T.H., Duboy, E.C. & Li, H. The Impact of DNA Topology and Guide Length on Target Selection by a Cytosine-Specific Cas9. ACS Synth Biol 6, 1103–1113 (2017).

21. Zheng, S.Q. et al. MotionCor2: anisotropic correction of beam-induced motion for improved cryo-electron microscopy. Nat Methods 14, 331–332 (2017).

22. Zhang, K. Gctf: Real-time CTF determination and correction. J Struct Biol 193, 1–12 (2016).

23. Punjani, A., Rubinstein, J.L., Fleet, D.J. & Brubaker, M.A. cryoSPARC: algorithms for rapid unsupervised cryo-EM structure determination. Nat Methods 14, 290–296 (2017).

24. Emsley, P. & Cowtan, K. Coot: model-building tools for molecular graphics. Acta Crystallogr D Biol Crystallogr 60, 2126–32 (2004).

25. Liebschner, D. et al. Macromolecular structure determination using X-rays, neutrons and electrons: recent developments in Phenix. Acta Crystallographica Section D-Structural Biology 75, 861–877 (2019).

26. Hand, T.H., Das, A. & Li, H. Directed evolution studies of a thermophilic Type II-C Cas9. Methods Enzymol 616, 265–288 (2019).

27. Hand, T.H. et al. Phosphate Lock Residues of Acidothermus cellulolyticus Cas9 Are Critical to Its Substrate Specificity. ACS Synth Biol 7, 2908–2917 (2018).

