## Supplementary Figures and Tables for "Conformational Changes Regulate Metal Coordination in the Catalytic Sites of Cas9"

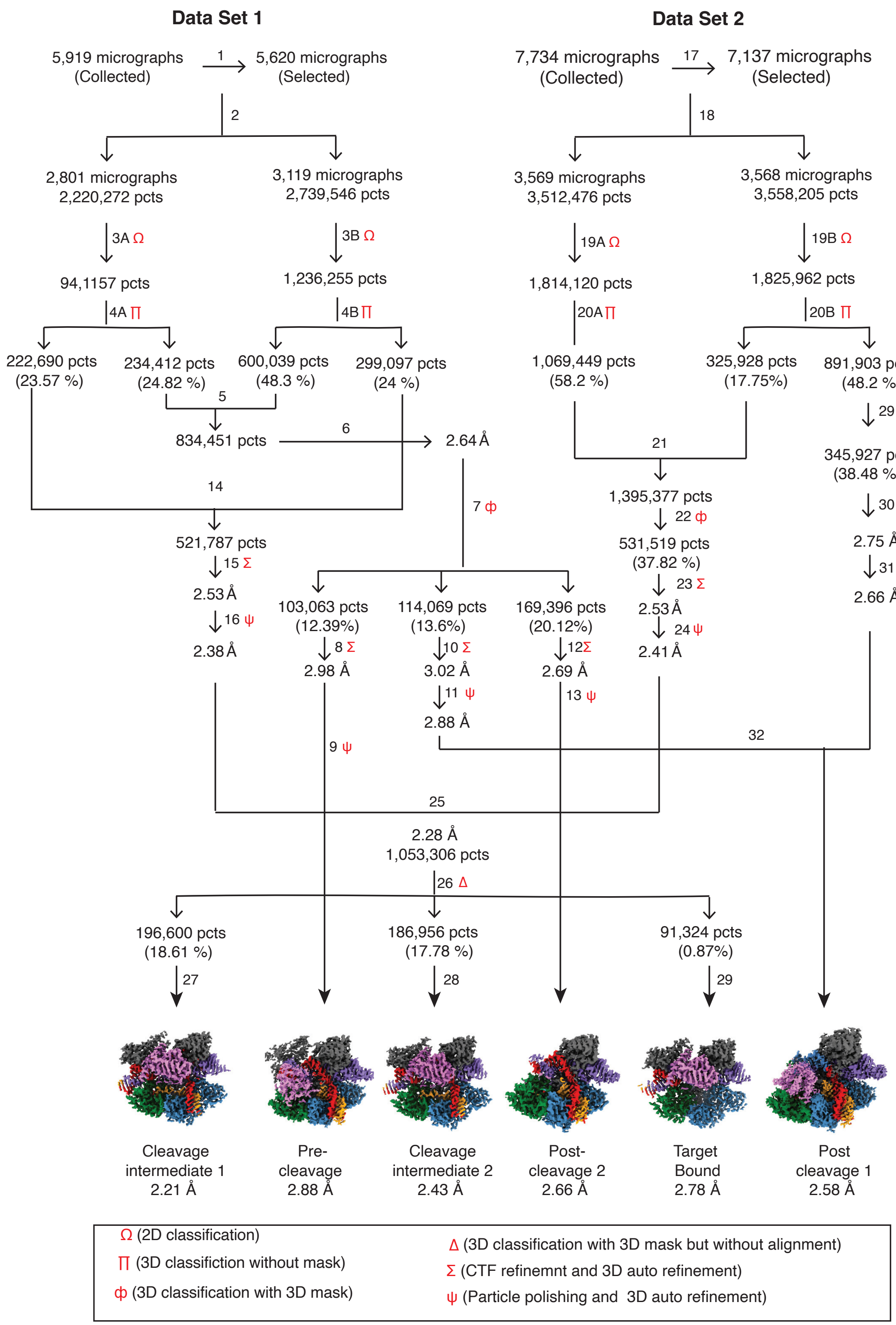


Supplementary Figure 1

**Supplementary Figure 1.** Workflow of single particle reconstruction of the active AceCas9 ternary complexes. Symbols in red represent the type of processes outlined in the textbox below. Numbers next to arrows account the steps of reconstruction and the percentage indicate those of particles to the steps above.

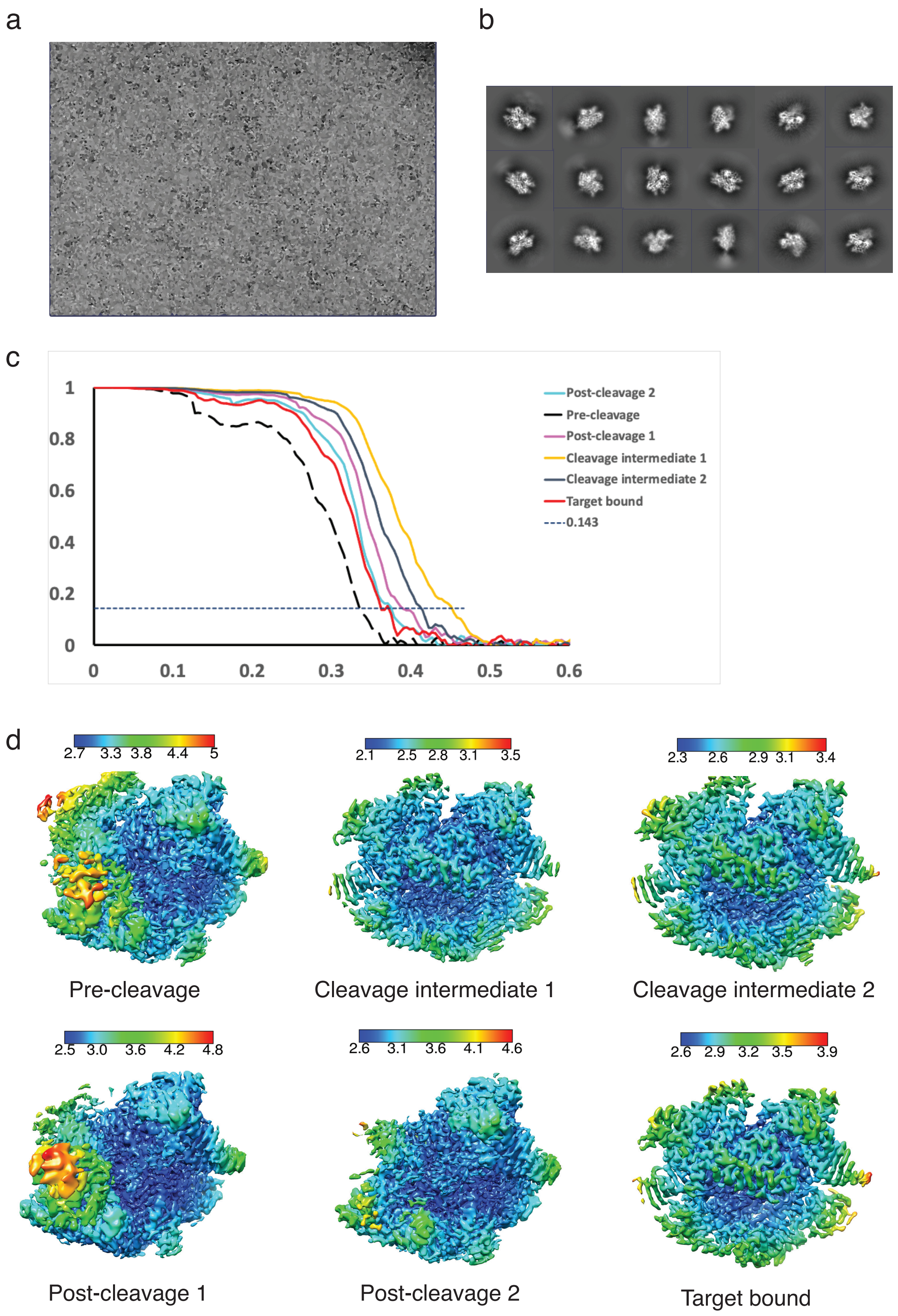


Supplementary Figure 2

**Supplementary Figure 2.** Map assessment of the cryoEM reconstructions. (a) Representative micrograph of the active AceCas9 ternary complex. (b) Select 2D class averages obtained from all particles used in reconstruction. (c) Fourier Shell Correlation (FSC) curves of the final classes that are colored and labeled, respectively. (d) Local resolution of the final six refined maps. The color bars define the resolution range.

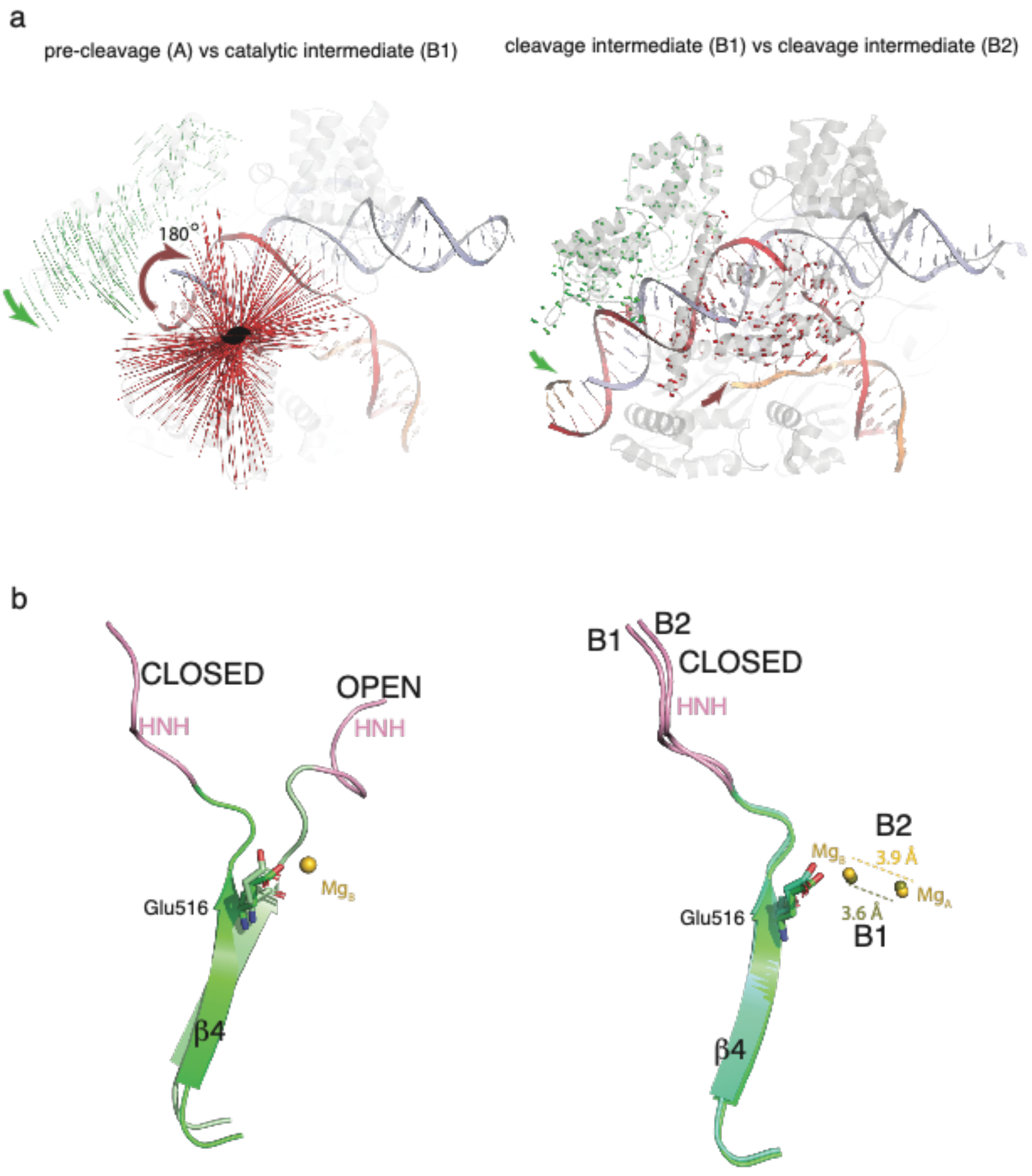


Supplementary Figure 3

**Supplementary Figure 3.** Conformational changes in AceCas9. (a) Comparison of domains when the sgRNA between the two compared complexes are aligned. Red lines indicate pair-wise displacement of Cα atoms within the HNH domain of the two compared AceCas9 structures. Green lines indicate pair-wise displacement of Cα atoms within the REC2 domain of the two compared AceCas9 structures. Curved arrows indicate the overall rotations of the HNH (red) and REC2 (green) domains. A 2-fold rotation symbol indicates the location of the axis of 2-fold rotation of the HNH domain. (b) Comparison of the β4 strand conformations and the location of Glu516 for the same pairs of compared structures in (a). Right, the dashed lines indicate the distance between the two metal ions in the two different conformations as labeled.

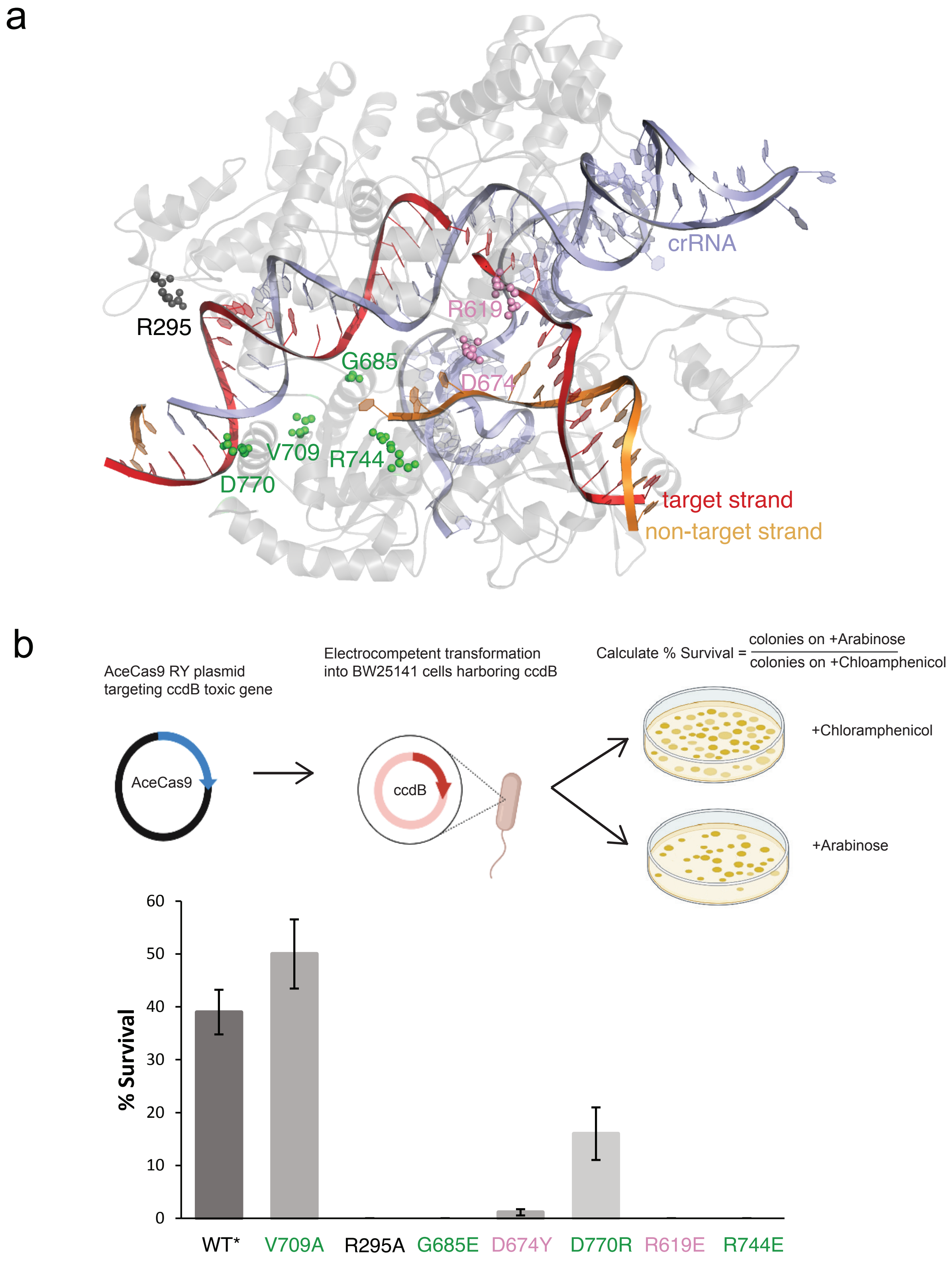


Supplementary Figure 4

**Supplementary Figure 4.** Assessment of enzyme activities of residues identified important for engaging the guide RNA-spacer heteroduplex. (a) Locations of the residues in the AceCas9 ternary complex shown in colored spheres and labeled. The same color scheme as in Figures 1 and 2 is used. (b) Top, schematic of the enzyme activity measurement by a modified cell survival assay (Materials and Methods). Bottom, percent of survival for the wild-type and the variants. WT* indicates AceCas9(Glu839Arg/Glu840Tyr) variant. Other mutations were constructed on the AceCas9(Glu839Arg/Glu840Tyr) variant.

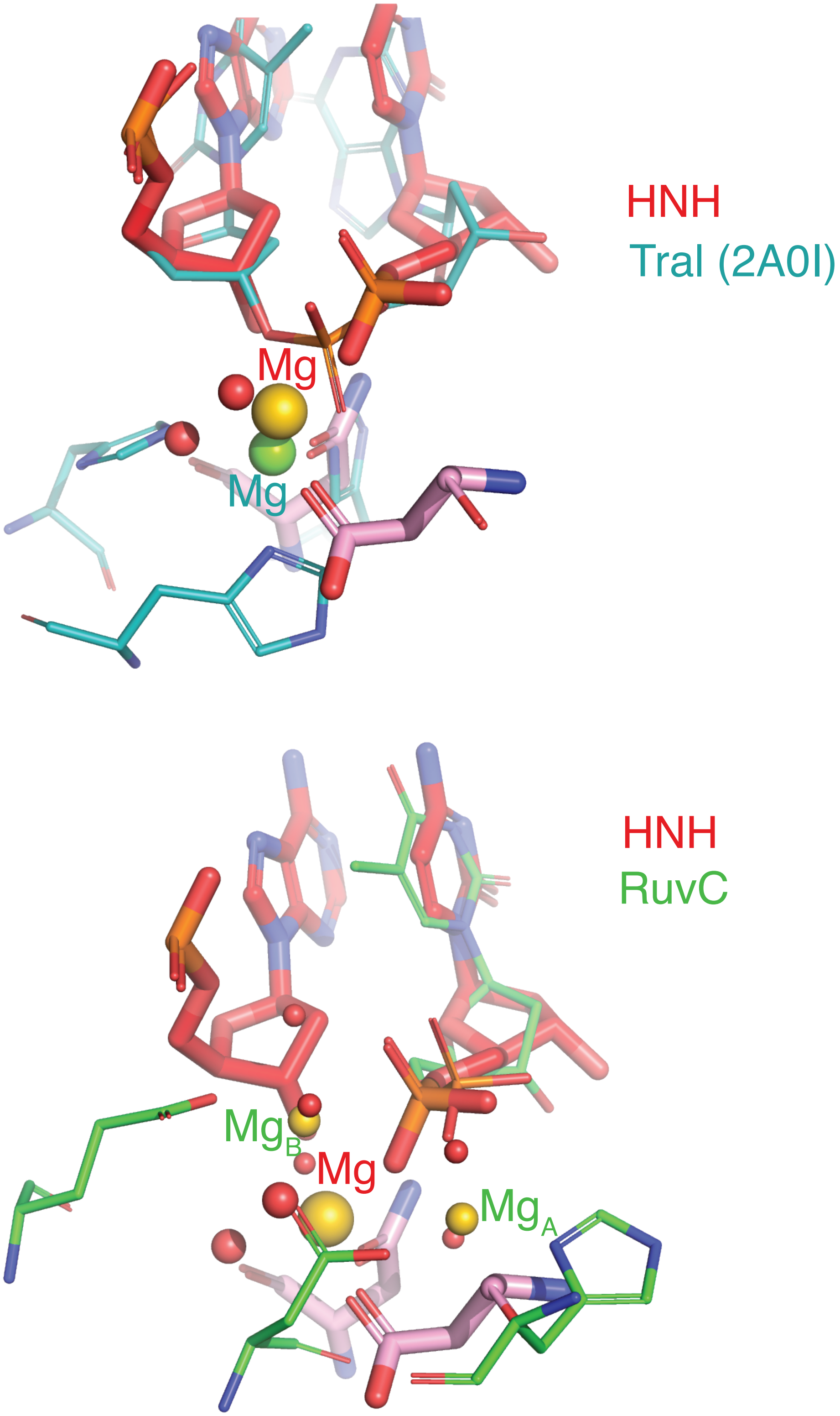


Supplementary Figure 5

**Supplementary Figure 5.** Comparison of active site geometry between HNH and TraI relaxase (PDB ID: 2A0I) (top) and between HNH and RuvC (bottom). Residues of TraI are shown as think teal sticks and the bound metal is drawn as a green sphere. Residues of HNH are shown as thick red sticks and the bound metal is drawn as a gold sphere. Residues of RuvC are shown as thin green sticks, metals are shown as small gold spheres, and water are shown as small red spheres.

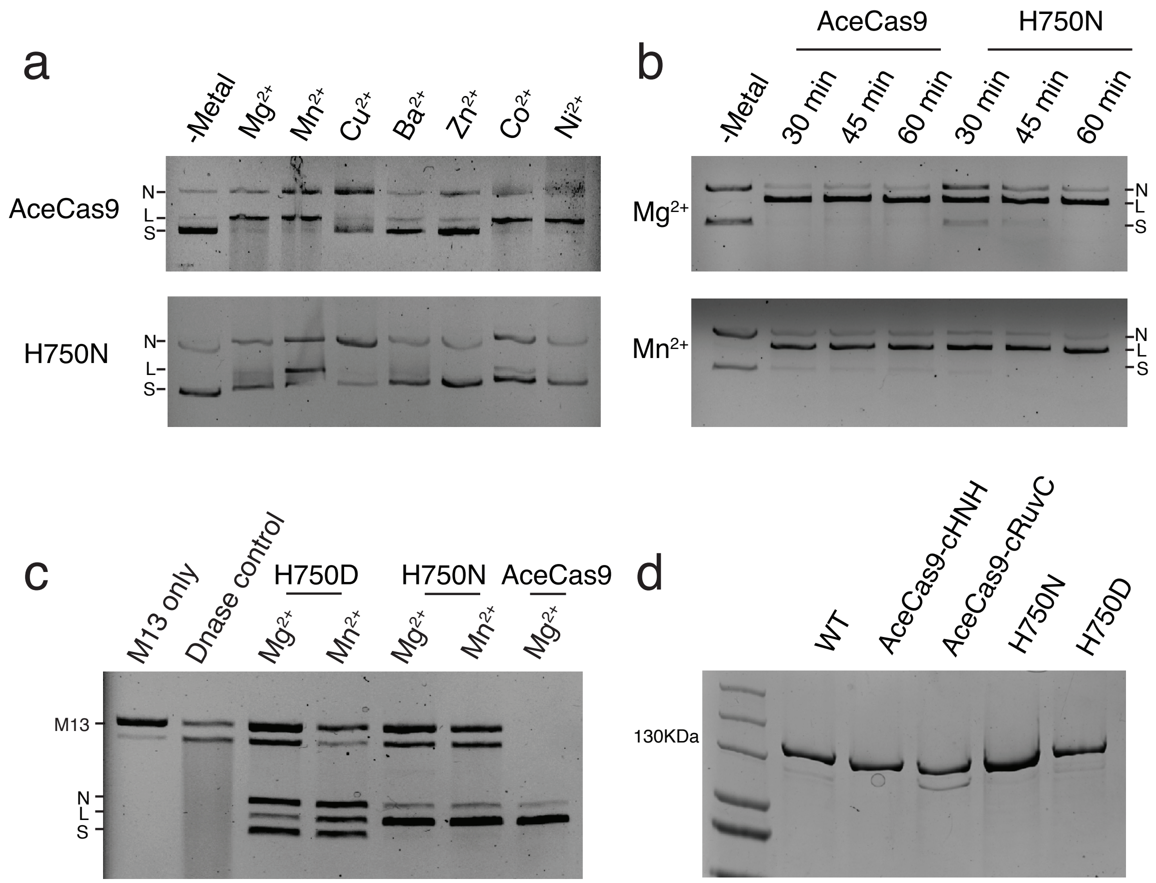


**Supplementary Figure 6.** Plasmid cleavage assay with AceCas9 wild type and mutants. (a) Target plasmid cleavage activities of AceCas9 and His750N with different divalent metals. "N" denotes nicked, "L" denotes linearized, and "S" denotes supercoiled plasmid DNA. (b) Comparison of DNA plasmid cleavage activities of AceCas9 and H750N with $Mg^{2+}$ and $Mn^{2+}$ at different timepoint. (c) M13 ssDNA cleavage (collateral activity) of AceCas9, H750D and H750N in presence of $Mg^{2+}$ or $Mn^{2+}$. Turbo DNase cleavage of M13 is used as the Dnase positive control. (d) Coomassie Blue stained SDS-PAGE of the proteins used in this study.

Pair-wise amino acid sequence identity of the known Cas9 (%) for RuvC/HNH

| | AceCas9 | AnaCas9 | Nme1Cas9 | SpyCas9 | SauCas9 | CdiCas9 | CjeCas9 |
|---|---|---|---|---|---|---|---|
| **AceCas9** | 100.0 | 22.3/27.9 | 16.8/16.7 | 13.2/12.5 | 14.5/15.9 | 22.7 | 15.9 |
| **AnaCas9** | | 100.0 | 14.8/17.5 | 15.6/11.8 | 12.6/12.6 | 36.0 | 13.9 |
| **Nme1Cas9** | | | 100.0 | 15.3/24.6 | 26.1/33.3 | 18.6 | 24.8 |
| **SpyCas9** | | | | 100.0 | 16.8/21.9 | 14.6 | 16.4 |
| **SauCas9** | | | | | 100.0 | 14.9 | 21.0 |
| **CdiCas9** | | | | | | 100.0 | 14.7 |
| **CjeCas9** | | | | | | | 100.0 |

HNH domain

```
Nme1Cas9HNH     ------------------------------------YFPNFVGEPKSKDILKLRLYEQQHGK
SauCas9HNH      ---------------------------------------TTGKENAKYLIEKIKLHDMQEGK
SpyCas9HNH      IEMARENQTTQKGQKNSRERMKRIEEGIKELGSQILKEHPVENTQLQNEKLYLYYLQNGR
AceCas9HNH      ------ASESRERQAEEEAARRAHRKANDRIRAELRASGLSDPSPADLVRARLLELYDCH
AnaCas9HNH      -------SERMADERDKANRRRYNDNQEAMKKIQRDYGKEGYISRGDIVRLDALELQGCA
                                                                 :

Nme1Cas9HNH     CLYSGKEINLG-RLNEKGYVEIDHALPFSRTWDD-SFNNKVLVLGSENQNKGNQTPYEYF
SauCas9HNH      CLYSLEAIPLEDLLNNPFNYEVDHIIPRSVSFDN-SFNNKVLVKQEENSKKGNRTPFQYL
SpyCas9HNH      DMYVDQELDIN----RLSDYDVDHIVPQSFLKDD-SIDNKVLTRSDKNRGKSDNVPSEEV
AceCas9HNH      CMYCGAPISWE-------NSELDHIVPRTDGGSN-RHENLAITCGACNKEKG-RRPFASW
AnaCas9HNH      CLYCGTTIGYH-------TCQLDHIVPQAGPGSNNRRGNLVAVCERCNRSKS-NTPFAVW
                 :*    :            ::** :* :   .:    * . .    *  *. . *

Nme1Cas9HNH     NGKD--NSREWQEFKARVE--------------TSRFPRSKKQRIL------------
SauCas9HNH      SSSD--SKISYETFKKHILNL---------AKGKGRISKTKKEYLLEE----------
SpyCas9HNH      VKKM--KNYWRQLLNAKLITQRKF--DNLTKAERGGLSELDKAGFIKRQ---------
AceCas9HNH      AETS--NRVQLRDVIDRVQKLKYSGNMYWTRDEFSRYKKSVVARLKRRTSDPEVIQSI
AnaCas9HNH      AQKCGIPHVGVKEAIGRVRGWRKQT-PNTSSEDLTRLKKEVIARLRRTQEDPEIDER-
                  .        .    ::                    .     :
```

RuvC domain

```
CdiCas9RuvC     --------MKYHVGIDVGTFSVGLAAIEVDDAGMPIKTLSLV---SHIHDSGLDPDKIKS
AnaCas9RuvC     -WYASLSAHHLRVGIDVGTHSVGLATLRVDDHGTPIELLSAL---SHIHDSGVGKEGKKD
AceCas9RuvC     ----GTVPVTWRLGVDVGERSIGLAAVSYEED-KPKEILAAV---SWIHDGGVALH----
Nme1Cas9RuvC    MAAFKPNSINYILGLDIGIASVGWAMVEIDEEENPIRLIDLG---VRVFERAEVPKTGKK
SauCas9RuvC     ------MKRNYILGLDIGITSVGYGIIDYETR----DVIDAG---VRLFKEANVEIPTTL
CjeCas9RuvC     --------MARILAFDIGISSIGWAFSENDELKDCGVRIFTK---AENPKTGESLAINED
SpyCas9RuvC     ------MDKKYSIGLDIGTNSVGWAVITDEYKVPSKKFKVLGNTDRHSIKKNLIGALLFD
                            :..*:*  *:* .    :                   .

CdiCas9RuvC     AVTIEPSWTPPAPRIGEPVGNPAVDRVLKTVSRWLESATKTWG--APERVIIEHV-----
AnaCas9RuvC     HDTRKKAIN-------APVGNPSVDRTLKIVGRYLS-AVESWG--TPEVIHVEHVRDGFT
AceCas9RuvC     ----------------EATGHPVVDRNLAILRKFLSSATMRWG--PPQSIVVELY-----
Nme1Cas9RuvC    NTEEKIYLPPIP---ADEIRNPVVLRALSQARKVINGVVRRYG--SPARIHIETAR----
SauCas9RuvC     VD--------------DFILSPVVKRSFIQSIKVINAIIKKYG--LPNDIIIELAR----
CjeCas9RuvC     KKDFLPAFN--ETYYKDEVTNPVVLRAIKEYRKVLNALLKKYG--KVHKINIELAR----
SpyCas9RuvC     SGETAEATDSLHEHIANLAGSPAIKKGILQTVKVVDELVKVMGRHKPENIVIEMAR----
                                     * : : :    : :.      *      : :*

CdiCas9RuvC     SMESVAWMANELRSRVAQHFASH-GTTVRVYRGSLTAEARRASGISGK-----LEFLDGV
AnaCas9RuvC     SSESVAWAN-ELHHRIAAAYP----ETTVVYRGSITAAARKAAGIDSR-----INLIGEK
AceCas9RuvC     -------AAVALRDRLLSYGEKNGVAQVAVFRGGVTAEARRWLDISIERLFSRVAIFAQS
Nme1Cas9RuvC    -------EVGLNDTRYVNRFLCQFVADRMRLTGKGKKRVFASNGQITN----LLRGFWGL
SauCas9RuvC     -------------ERYATRGLMNLLRSYFRVNN-LDVKVKSINGGFTS----FLRRKWKF
CjeCas9RuvC     -YIARLVLNYTKDYLDFLPLSDDENTKLNDTQKGSKVHVEAKSGMLTS----ALRHTWGF
SpyCas9RuvC     ----------RQITKHVAQILDSRMNTKYDENDKLIREVKVITLKSKLVS--DFRKDFQF
                                                      .              .

CdiCas9RuvC     G-KSRLDRRHHAIDAAVIAFTSDYVAETLAVRSNLKQSQAHR-----------QEAPQWR
AnaCas9RuvC     GRKDRIDRRHHAVDASVVALEAS-VAKTLAERSSLRGEQRLT-----------GKEQTWK
AceCas9RuvC     TSTKRLDRRHHAVDAVVLTTLTPGVAKTLADARSRRVSASTE-----------EPQ----
Nme1Cas9RuvC    RKVRAENDRHHALDAVVVACSTVAMQQKITRFVRYKEMNAFDGKTIDKETGEVLHQKTHF
SauCas9RuvC     KKERNKGYKHHAEDALIIANADFIFKEWKKLDKAKKVMENQMFE----------------
CjeCas9RuvC     STKDRNNHLHHAIDAVIIAYANNSIVKAFSDFKKEQESNSAELY----------------
SpyCas9RuvC     YKVREINNYHHAHDAYLNAVVGTALIKKYPKLESEFVYGDYKVYDVRKMIAK-SEQEIGK
                      .  *** ** : :    . :

CdiCas9RuvC     EFTGKDAEHRAAWRVWCQKMEKLSALLTEDLRDDRVVVMSNVR-----------------
AnaCas9RuvC     QYTGSTVGAREHFEWRG--HLHLTELFNERLAEDKVYVTQNIRLRLSD------------
AceCas9RuvC     ---------SPAYRQWKESCSGLGDLLISTAARDSIAVAAPLRLRP--------------
Nme1Cas9RuvC    PQPWEFFAQEVMIRVFGKPDGKPEFEEADTLEKLRTLLAEKLSSRPEAVHEYVTPLFVSR
SauCas9RuvC     -----EKQAESMPEIETEQEYKEIFITPHQIKHIKDFKDYKYSHRV--------------
CjeCas9RuvC     --------AKKISELDYKNKRKFFEPFSGFRQKVLDKIDEIFVSKP--------------
SpyCas9RuvC     ATAKYFFYSNIMNFFKTEITLANGEIRKRPLIETNGETGEIVWDKGRDFATVRKVLSMPQ
```

Supplementary Figure 7

**Supplementary Figure 7.** Comparison of sequences of the RuvC and HNH domains for the known Cas9s. Sequences are extracted from the domains of the three-dimensional structures of the Cas9. Top, pair-wise sequence identities for the known Cas9. Bottom, sequence alignment for the HNH (top) and the RuvC domain (bottom), respectively. The strictly conserved catalytic residues are highlighted by red outlines.

**Table S1a:** Statistics of cryo-EM data collection processing

| **Data acquisition and processing parameters** | **Data-set 1** | **Data-set 2** |
|---|---|---|
| Microscope | Titan Krios G3i | Titan Krios G3i |
| Detector | Gatan K3 | Gatan K3 |
| Voltage | 300 kV | 300 kV |
| Collecting mode | Counted super-resolution | Counted super-resolution |
| Dose rate ($e^-$/ $Å^2$) | 60 | 60 |
| Defocus range (μm) | (-1) - (-2.2) | (-1) - (-2.2) |
| Nominal magnification | 105K | 105K |
| Frames collected per exposure | 60 | 60 |
| Frame-alignment software | MotionCor2 | MotionCor2 |
| CTF estimation software | Gctf | Gctf |
| Raw images collected | 5919 | 7734 |
| Images used for particle picking | 5620 | 7137 |
| 2D classification software | Cryosparc | Cryosparc |
| Final reconstruction software | RELION-4 | RELION-4 |
| Applied symmetry | C1 | C1 |
| Resolution method | FSC 0.143 cutoff | FSC 0.143 cutoff |
| Local resolution software | RELION-4 | RELION-4 |
| Map visualization software | Pymol/Chimera/Chimera-X, Coot | Pymol/Chimera/Chimera-X, Coot |

**Table S1b:** Statistics of model refinement and data deposition

| **Refinement parameters** | **Pre-cleavage** | **Cleavage-intermediate 1** | **Cleavage-intermediate 2** | **Post-cleavage 1** | **Post-cleavage 2** | **Target bound** |
|---|---|---|---|---|---|---|
| Deposited EMDB | **27143** | **27142** | **27141** | **27146** | **27144** | **27145** |
| CC (mask) | 0.73 | 0.87 | 0.84 | 0.78 | 0.83 | 0.82 |
| RMSD (Bond lengths/Bond angles) | 0.006/0.953 | 0.007/0.793 | 0.006/0.741 | 0.005/0.834 | 0.004/0.820 | 0.005/0.808 |
| Number of particles contributed to the final reconstruction | 103,063 | 196,600 | 186,956 | 459,996 | 169,296 | 91,324 |
| Final resolution (Å) | 2.88 | 2.21 | 2.43 | 2.58 | 2.66 | 2.78 |
| Ramachandran plot (Outliers<br>Allowed<br>Favored) | 0.00<br>4.93<br>95.07 | 0.00<br>1.82<br>98.18 | 0.00<br>1.46<br>98.54 | 0.00<br>3.34<br>96.55 | 0.14<br>2.30<br>97.56 | 0.00<br>2.1<br>97.90 |
| Cβ Outliers (%) | 0.00 | 0.10 | 0.00 | 0.00 | 0.00 | 0.00 |
| ADP (B-factors)<br>Iso/Aniso (#)<br>Protein<br>Nucleotide<br>Ligand<br>Water<br>(min/mask/mean) | 11415/0<br>0.0/52.7/19.2<br>0.0/68.2/24.2<br>17.8/17.8/17.8<br>--- | 12228/0<br>0.0/61.9/24.6<br>0.0/119.0/40.3<br>14.5/63.0/47.1<br>2.2/30.81/18.1 | 12295/0<br>0.0/63.7/27.8<br>0.0/127.2/44.7<br>13.8/79.4/60.7<br>2.6/48.6/20.2 | 9722/0<br>0.00103.2/38.2<br>1.7/186.4/70.6<br>15.8/150.9/69.6<br>24.4/32.6/28.5 | 8472/0<br>0.0/71.2/34.1<br>6.1/123.1/57.1<br>24.4/56.3/34.5<br>21.6/41.8/33.8 | 11436/0<br>0.0/59.4/25.2<br>0.0/116.8/37.1<br>6.6/69.0/54.2<br>0.0/29.3/14.7 |
| MolProbity score | 1.65 | 1.38 | 1.45 | 1.44 | 1.25 | 1.16 |
| Clash score | 5.68 | 6.85 | 8.36 | 4.34 | 3.71 | 3.50 |
| Rotamer outlier (%) | 0.00 | 0.00 | 0.00 | 0.00 | 0.00 | 0.00 |
| FSC model (0/0.143/0.5) | 2.4/2.5/3.1 | 2.0/2.0/2.3 | 2.0/2.1/2.5 | 2.0/2.2/2.7 | 2.2/2.3/2.8 | 2.2/2.3/2.8 |
| Deposited PDB codes | **8D2N** | **8D2L** | **8D2K** | **8D2Q** | **8D2O** | **8D2P** |

**Supplementary Table 2.** DNA or RNA Oligos used for this study

| Name | Sequence (5'-3') | Used for |
|---|---|---|
| **40mer NTS**<br>**40mer TS** | TCTAGAGGTAGGATGGCAAGATCCTGGTATACACCAAGCT<br>AGCTTGGTGTATACCAGGATCTTGCCATCCTACCTCTAGA | Cryo EM |
| **DNA for sgRNA106** | GAA CCC CCT CGC TGC TGC GAG GGGGTG AAG AAT GCG ACC CCA CGA AGGGGT CTT GCT AGG TAG CCT TTT CAGGCT CCC CAG CAT ACC AGG ATC TTGCCA TCC TAC CTA TAG TGA GTC GTATTA | Cryo EM |
| **sgRNA106** | GGUAGGAUGGCAAGAUCCUGGUAUGCUGGGGAGCCUGAAAAGGCUACCU AGCAAGACCCCUUCGUGGGGUCGCAUUCUUCACCCCCUCGCAGCAGCGAG GGGGUUC | Cryo EM |
| **FAM NTS**<br>**TS** | [FAM] GGT AGG ATG GCA AGA TCC TGGTAT ACA CCA AGC T<br>AGC TTG GTG TAT ACC AGG ATC TTGCCA TCC TAC C | Oligo Cleavage assay |
| **NTS**<br>**HEX TS** | TCTAGAGGTAGGATGGCAAGATCCTGGTATACACCAAGCT<br>[HEX] AGCTTGGTGTATACCAGGATCTTGCCATCCTACCTCTAGA | Oligo Cleavage assay |
| **G685E-F**<br>**G685E-R** | GAATCCACCGAATACGCAGCTG<br>AATGCTTTGAATCACTTCGGG | Q5 site-directed mutagenesis |
| **V709E-F**<br>**V709E-R** | CAGGTAGCGGAATTCCGCGGTG<br>AGCTACTCCGTTCTTTTCGCCG | Q5 site-directed mutagenesis |
| **D674Y-F**<br>**D674R-F**<br>**D674Y_R-R** | CGCACCTCCTATCCCGAAGTG<br>CGCACCTCCCGTCCCGAAGTG<br>CCTTTTGAGGCGGGCGACGAC | Q5 site-directed mutagenesis |
| **D770R-F**<br>**D770R-R** | GAAGACCTTGGCCAGAGCGCGGAGTC<br>GCGACGCCCGGAGTCA | Q5 site-directed mutagenesis |
| **R295A-F**<br>**R295A-R** | GCTAACCTGGCGATACGTGATGG<br>CACCGCCGCAACGATGCGATAC | Q5 site-directed mutagenesis |
| **R744E_F**<br>**R744E_R** | GCACGAAGGAACTCGATCGTCGGC<br>TCGTTGACTGAGCAAAAATTGC | Q5 site-directed mutagenesis |
| **R619E_F**<br>**R619E_R** | GGA AAA AGG TGA ACG TCC CTT TGCG<br>TTG TTG CAC GCT CCG CAG G | Q5 site-directed mutagenesis |
| **H750N_F**<br>**H750n_R** | CGT CGG CAC AAC GCC GTG GAC GCG<br>ATC GAG CCG CTT CGT GCT CGT TGA C | Q5 site-directed mutagenesis |
| **H750D_F**<br>**H750D_R** | GTC GGC ACG ATG CCG TGG ACG CG<br>GAT CGA GCC GCT TCG TGC TCG TTG | Q5 site-directed mutagenesis |
